# Macula densa-targeting gene delivery in vivo using AAV9

**DOI:** 10.64898/2026.11.15.766863

**Authors:** Georgina Gyarmati, Alejandra Becerra Calderon, Audrey Izuhara, Vadym Buncha, Sachin Deepak, Karam Aboudehen, János Peti-Peterdi

## Abstract

Macula densa (MD) cells are key sensory and regulatory cells in the kidney; however, they have been difficult to study because of a lack of cell-specific, user-friendly in vivo research tools. Here we demonstrate the high affinity of wild-type Adeno-Associated Virus serotype 9 (AAV9) for MD cells and the feasibility of AAV9-mediated MD cell-specific genetic manipulations when administered to neonatal mice or after a temporary physiological challenge (low-salt diet) in the adult. A single intraperitoneal injection of low-dose AAV9-GFP given to neonatal mice on postnatal days P2-3 resulted in highly MD-specific and efficient transduction (∼85% of all GFP+ cells were Nos1+ MD cells). Proof-of-concept studies used AAV9 to overexpress or knock down biologically relevant genes in MD cells and establish MD gain- and loss-of-function (Gof/Lof) mouse models for initial functional and structural phenotyping using intravital multiphoton microscopy at the single-nephron level and immunofluorescence-based histological analysis. RNAscope confirmed the MD-specific knockdown of Pregnancy-associated plasma protein A2 (Pappa2) expression by AAV9-Pappa2 shRNA treatment in MD-Pappa2^Lof^ mice, and the MD- specific overexpression of Cell communication network 1 (Ccn1) by AAV9-Ccn1 mRNA treatment in MD-Ccn1^Gof^ mice. The mouse phenotypes revealed alterations in classic MD/juxtaglomerular functions (renin control and tubuloglomerular feedback-mediated glomerular hemodynamics) as well as in newly emerging MD cell functions (tissue remodeling). MD-specific genetic manipulations in vivo via AAV9-mediated gene delivery may enable future studies of MD cell functions, including the MD program for endogenous kidney tissue remodeling and regeneration, and support the development of MD cell-targeting gene therapies for kidney diseases.

**NEW & NOTEWORTHY:** The AAV9 gene delivery approach described in this study may be useful for straightforward in vivo genetic manipulation of the minority renal cell type, the macula densa (MD), in preclinical renal physiology and disease research. This novel research tool may improve our understanding of the classic renal hemodynamic and renin-regulatory functions of the understudied MD cells, as well as their newly emerging roles in kidney tissue remodeling and regeneration and in the renal nervous system. In the long term, AAV9-mediated MD targeting gene delivery may enable the development of gene therapies for a range of kidney diseases.

## Introduction

Genetic manipulations that overexpress or knock down specific molecular targets, or replace or introduce genetic material in vivo, are highly specific, powerful, and widely used preclinical research strategies for studying protein and cell functions. Gene therapies are also rapidly emerging in the clinic to treat or cure chronic or inherited genetic diseases. Commonly used in vivo gene delivery methods include viral (adeno-associated viruses (AAVs), lentiviruses) and non-viral vectors (lipid nanoparticles (LNPs), polymeric nanoparticles, extracellular vesicles, liposomes), and physical methods (electroporation, hydrodynamic delivery)(1, 2). Common bottlenecks in gene delivery methods include limited cell- and tissue-specific tropism and payload-size constraints (2).

AAVs are commonly used vectors for in vivo gene transfer because of their favorable safety profile, low immunogenicity, and lack of integration into the host genome (2). Different naturally occurring or synthetic AAV serotypes (AAV2-9) allow tissue-specific tropism for different organs, including the liver, muscle, central nervous system, and eye, with AAV9 appearing most effective for kidney cell transduction (3–6). Depending on the route of administration and tissue conditions, previous studies found effective AAV9-mediated transduction of mesangial and interstitial cells, podocytes, and proximal tubules (3–7). However, AAV9 has tropism for other organs including the heart, skeletal muscle, and brain (2, 8, 9). Unlike other AAV serotypes, AAV9 can cross the blood-brain barrier and therefore has well-established tropism for neurons. In one of the first AAV-based human gene therapies, AAV9 was used to treat spinal muscular atrophy (10). Interestingly, and importantly for the present work, animal studies found that AAV9 can more efficiently target cells in the central nervous system, including neurons, if given before postnatal day 5 (9).

The macula densa (MD) is a group of 20-25 highly specialized renal epithelial cells in each nephron at the distal end of the thick ascending limb, strategically localized at the glomerular vascular pole as the tubular component of the juxtaglomerular apparatus (JGA). The classic MD cell functions include the control of renin synthesis and release from the adjacent JG renin cells (11, 12) and tubuloglomerular feedback (TGF)-mediated regulation of renal and glomerular hemodynamics, including renal blood flow (RBF) and glomerular filtration rate (GFR)(11, 13, 14). However, recent research suggests MD cells are neuronally differentiated and more complex than previously thought and play emerging roles in endogenous kidney tissue remodeling and regeneration, as well as in the renal nervous system (15–19). Recent high-resolution transcriptome analysis of mouse MD cells found enrichment for developing brain-like genes rather than kidney-tissue-specific genes (16).

Given the evidence of effective AAV9-mediated targeting of renal cells and neurons, the neuronal characteristics of MD cells, and the fact that embryonic developmental conditions boost neuronal targeting, we hypothesized that AAV9 may have a high affinity for MD cells in neonatal mice and under certain adult stress conditions that recapitulate developmental states. One such condition is low dietary salt intake and ACE inhibitor treatment, which result in loss of body fluid and salt (16, 20). Therefore, the present study aimed to test AAV9-mediated gene delivery targeting MD cells and provide proof of concept for the feasibility of MD-targeting genetic manipulations in vivo.

## Materials and Methods

### Animals and Treatments

All animal protocols were approved by the Institutional Animal Care and Use Committee at the University of Southern California. Male and female C57BL6/J mice were used from our breeding colony at USC either at postnatal days P2-3 or at 8-10 weeks old (originally purchased from Jackson Laboratory, Bar Harbor, ME). For afferent arteriole vascular smooth muscle cell calcium imaging studies, Ren1d-GCaMP6f/tdTomato (Salsa6f) mice were used, which were generated by crossing Ren1d-Cre (20) and Salsa6f floxed (RRID:IMSR_JAX:031968)(21) mice resulting in the expression of the ratiometric fusion calcium indicator Salsa6f (tdTomato linked to GCaMP6f by a V5 epitope tag) in cells of the renin lineage as described before (22). Some mice received a low-salt diet for 10 days (TD 90228, Harlan Teklad, Madison, WI).

To generate mice with MD-specific knockdown of Pappa2 (MD-Pappa2^Lof^), mice were injected intraperitoneally (ip.) with a single low-dose of mouse Pappa2 shRNA and EGFP marker containing pAAV[shRNA]-CAG>EGFP-U6>mPappa2 vector (AAV9, 10^11^ GC in 50 μL saline, Vector ID: VB900174-8501wdk, VectorBuilder, Chicago, IL)(23, 24). For the generation of mice with MD-specific overexpression of Ccn1 (MD-Ccn1^Gof^), animals were injected ip. with AAV9 virus containing pAAV[Exp]-CMV>mCcn1[NM_010516.2]:WPRE (AAV9, 10^11^ GC in 50 ul PBS, Vector ID: VB900169-7858zta). A scrambled sh/mRNA control AAV9 virus construct (Vector ID: VB010000-9489hhg) expressing GFP in target cells was used as a negative control (24).

### Immunohistochemistry and RNAscope of fixed kidney tissue

Immunofluorescence detection of proteins was performed as described previously (25). Briefly, kidneys were perfused and fixed in 4% PFA for 2 hours at room temperature, embedded in paraffin, and sectioned 8 µm thick. For antibody stains, slides were washed in 1XPBS. For antigen retrieval, heat-induced epitope retrieval was applied by boiling sections in Sodium Citrate buffer (pH 6.0) or Tris-EDTA (pH 9.0) for ten minutes. To reduce non-specific binding, sections were blocked with normal goat serum (1:20). Primary and secondary antibodies were applied sequentially overnight at 4° C and 2 hours at room temperature. Primary antibodies and dilutions were as follows: Nos1 (Santa Cruz Biotechnology, Santa Cruz, CA, 1:100), PDGFRb (3169S, Cell Signaling Technology, Danvers, MA, 1:100), renin (AS-54371, AnaSpec Inc 1:100), WT1 (ab89901, Abcam, 1:100), CD31 (No. 77699; Cell Signaling Technology, Danvers, MA, 1:100), GFP (GFP917979, Aves Labs,1:200). Alexa Fluor 488 and 594-conjugated secondary antibodies were purchased from Thermo Fisher Scientific and Jackson ImmunoResearch Laboratories. Slides were mounted by using DAPI-containing mounting media (VectaShield, Vector Laboratories Inc., Burlingame, CA).

Pappa2 and Ccn1 mRNA detection combined with GFP immunostaining in mouse kidney sections were manually carried out using the RNAscope Multiplex Fluorescent v2 Assay with TSA Vivid Dyes (323 280; Advanced Cell Diagnostics, Newark, CA), and the RNA-Protein Co-Detection Ancillary Kit (323 180; Advanced Cell Diagnostics) according to manufacturer’s instructions. Briefly, 4 μm formalin-fixed, paraffin-embedded slides were used. RNAscope Co-Detection Target Retrieval reagent was used for antigen-retrieval. Primary (GFP, GFP917979, Aves Labs, 1:200) and secondary antibodies (703-545-155, Jackson ImmunoResearch Laboratories), and mRNA Probe- Mm-Pappa2 and Ccn1 (Advanced Cell Diagnostics) and RNAscope Multiplex Fluorescent Detection Reagents were applied sequentially.

For structural analysis, high spatial resolution Z-stack recordings of identical tissue volumes and maximum intensity projection images (MIP) were used. Individual cell counting was performed based on immunofluorescence labeling with cell-specific markers and DAPI labeling of cell nuclei, as described before (16, 22). Only those cortical nephrons were selected in the tissue sections that contained the mid-sections of glomeruli, in which both the vascular pole (MD cells) and the glomerulo-tubular junction were visible. The sections were identified by animal number only rather than by experimental group, which was unknown to the examiner. Sections were examined with a Leica Stellaris WLL Falcon (Leica Microsystems, Wetzlar, Germany) confocal/multiphoton laser scanning microscope system. Imaging software (Leica LAS X and Image J, National Institutes of Health and the Laboratory for Optical and Computational Instrumentation (LOCI, University of Wisconsin) was used to measure glomerular tuft area and to calculate signal density as described before (16).

### Multiphoton microscopy (MPM)

Mice were anesthetized with isoflurane (1-3%) and prepared for intravital imaging of the intact living kidney, performed using a Leica Stellaris DIVE WLL multiphoton microscope (MPM) (Leica Microsystems, Wetzlar, Germany) powered by a Discovery NX laser system (Coherent Inc., Santa Clara, CA) as described previously (26–28). Images were acquired using a Leica 25× water-immersion objective (numerical aperture (NA) 1.1) and excitation at either 860 nm or 970 nm and a DMI8 inverted microscope’s external Leica 4Tune spectral hybrid detectors (emission at, 510-530 nm for GFP and GCaMP6, 580-600 nm for tdTomato). We minimized potential toxicity from laser excitation and fluorescence by using low laser power and high scan speeds to keep total laser exposure as low as possible. Image acquisition (12-bit, 512×512 pixel) consisted of only one z stack or time series per tissue volume (<2-10 min), which resulted in no apparent cell injury.

To study dynamic changes in intracellular calcium signaling in afferent arteriole vascular smooth muscle cells, fluorescence images were collected in time series (xyt, 526 ms per frame) with the Leica LAS X imaging software and using the same instrument settings (laser power, offset, gain of both detector channels) as described before (16). MPM imaging was always performed at the same time of day.

### Statistical methods

Data were expressed as average ± SEM and were analyzed using Student’s t-tests (between two groups). For illustration purposes, data were presented in box plots as a center line (mean), box (25th–75th percentiles), and whiskers (minimum to maximum values). Normality analysis was performed by using the Shapiro–Wilk test. P<0.05 was considered significant. There were no animals, data points, or experimental units excluded from the study. Statistical analyses were performed using GraphPad Prism 9.0.1 (GraphPad Software, Inc., La Jolla, CA).

## Results

### Macula densa-specific AAV9 tropism in the neonatal versus adult kidney

Based on the established tropism of wild-type AAV9 for the kidney (3) and its age-dependent cell targeting in other organs (9), our initial studies tested the efficacy of wild-type AAV9-mediated transduction in the adult and neonatal kidney. When adult mice were given a single systemic intraperitoneal (ip) injection of AAV9-GFP at various doses (10^11^-10^13^ GC in 50 ul PBS) and tissues harvested 2-4 weeks later for histological analysis, essentially no GFP-labeled cells were observed in the kidney. In striking contrast, when a low-dose injection of AAV9-GFP (10^11^ GC in 25 ul PBS ip.) was given to neonatal mice on postnatal days P2-3 and tissues harvested 4-6 weeks later, it resulted in strong GFP expression, which was limited to a few tubular epithelial cells at the glomerular vascular pole in all nephrons (Fig. 1A). Co-localization of GFP and the MD cell marker Nos1 immunolabeling identified the GFP+ cells as macula densa cells (Fig. 1B). The AAV9-mediated GFP expression was highly MD-specific, as approximately 85% of all GFP+ cells were also Nos1+ (Fig. 1C). In addition, MD cell transduction by AAV9 was also highly efficient, as approximately 80% of all Nos1+ cells were also GFP+ (Fig. 1D). Only rare and sporadic GFP labeling was found in the renal medulla, and the remaining approx. 15% GFP+ non-MD cells in the cortex were mesangial, renin, and endothelial cells (identified by PDGFRb, renin, and CD31 co-labeling, respectively) at or near the glomerular vascular poles (not shown). As expected, AAV9-mediated GFP expression at the neonatal stage was also detected in other organs, including the brain, liver, and heart, although at a moderate level with the applied low-dose AAV9-GFP (Fig. 1E). Using xyz volumetric imaging, AAV9-GFP expression was also detected in renal nerve fibers in both the adult and neonatal models, in contrast to MD cell labeling, which was restricted to the neonatal model (Fig. 1G).

**Figure 1.**
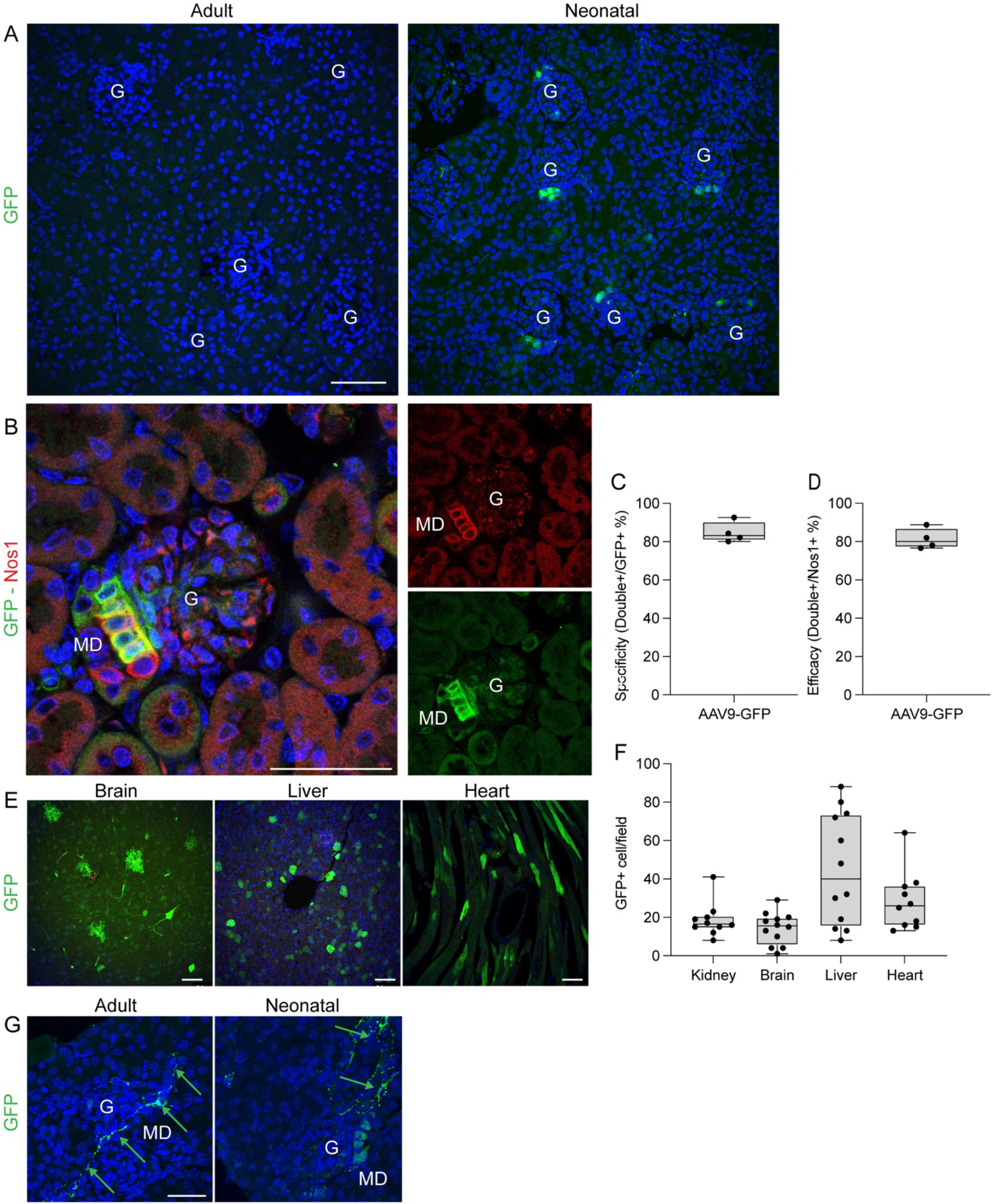
Histological features of the kidney and other organs after AAV9-GFP gene delivery in vivo in adult and neonatal conditions. A: Overview GFP immunofluorescence (green) images of large renal cortical areas with multiple glomeruli (G) from mice that were injected with AAV9-GFP at adult (left) or neonatal age (right). Cell nuclei were labeled blue with DAPI. Note the presence of several GFP+ cells at the vascular pole area of each glomerulus in the neonatal but not in the adult condition. **B:** Immunofluorescence double labeling of the macula densa (MD) cell marker Nos1 (red, top right) and the AAV9-driven genetic expression of GFP (green, bottom right) in a mouse kidney section. The overlay image (left) shows co-localization (yellow), confirming MD-specific GFP expression. **C-D:** Statistical summary of the MD specificity (C) and efficacy (D) of AAV9-driven gene delivery in mice. Data are presented in box plots as a center line (mean), box (25th–75th percentiles), and whiskers (minimum to maximum values). N=4 mice; data points represent the average of 10 nephrons each. **E:** GFP immunofluorescence images of brain, liver, and heart tissue sections from mice that were injected with AAV9-GFP at the neonatal age. **F:** Statistical summary of the number of GFP+ cells per field in each organ. N=10-12 mice each; data points represent the average of 3 different tissue areas in each. **G:** Xyz volumetric projection images of GFP immunofluorescence in the adult and neonatal models. Note the GFP+ renal nerve fibers in both conditions (arrows), but MD cells are GFP+ only in the neonatal condition. Scale bars are 50 µm.

### Low-salt diet reactivates juxtaglomerular tropism of AAV9 in the adult kidney

Stress conditions in the adult are known to temporarily reactivate embryonic developmental programs to facilitate repair and maintain tissue integrity. For example, the loss of body fluid and salt triggers a phenotypic change in multiple cells of the juxtaglomerular apparatus in the kidney to reestablish homeostasis (16, 20). Therefore, we next tested the effects of temporary dietary salt restriction on AAV9 tropism in the adult kidney. Adult mice were subjected to a low-salt diet for 10 days and then received a single ip. injection of AAV9-GFP (10^12^ GC in 50 ul PBS), and kidneys were harvested two weeks later. In contrast to the lack of GFP labeling in the untreated adult (Fig. 1A), the low-salt diet condition resulted in strong AAV9-mediated GFP expression in the kidney cortex, again limited to a few tubular epithelial cells at the glomerular vascular pole in all nephrons (Fig. 2, A and B). There was partial co-localization of GFP and Nos1 immunolabeling, suggesting that the GFP+ cells are in part MD cells, and in part “perimacular” cells of the thick ascending limb-distal tubule (Fig. 2C).

**Figure 2.**
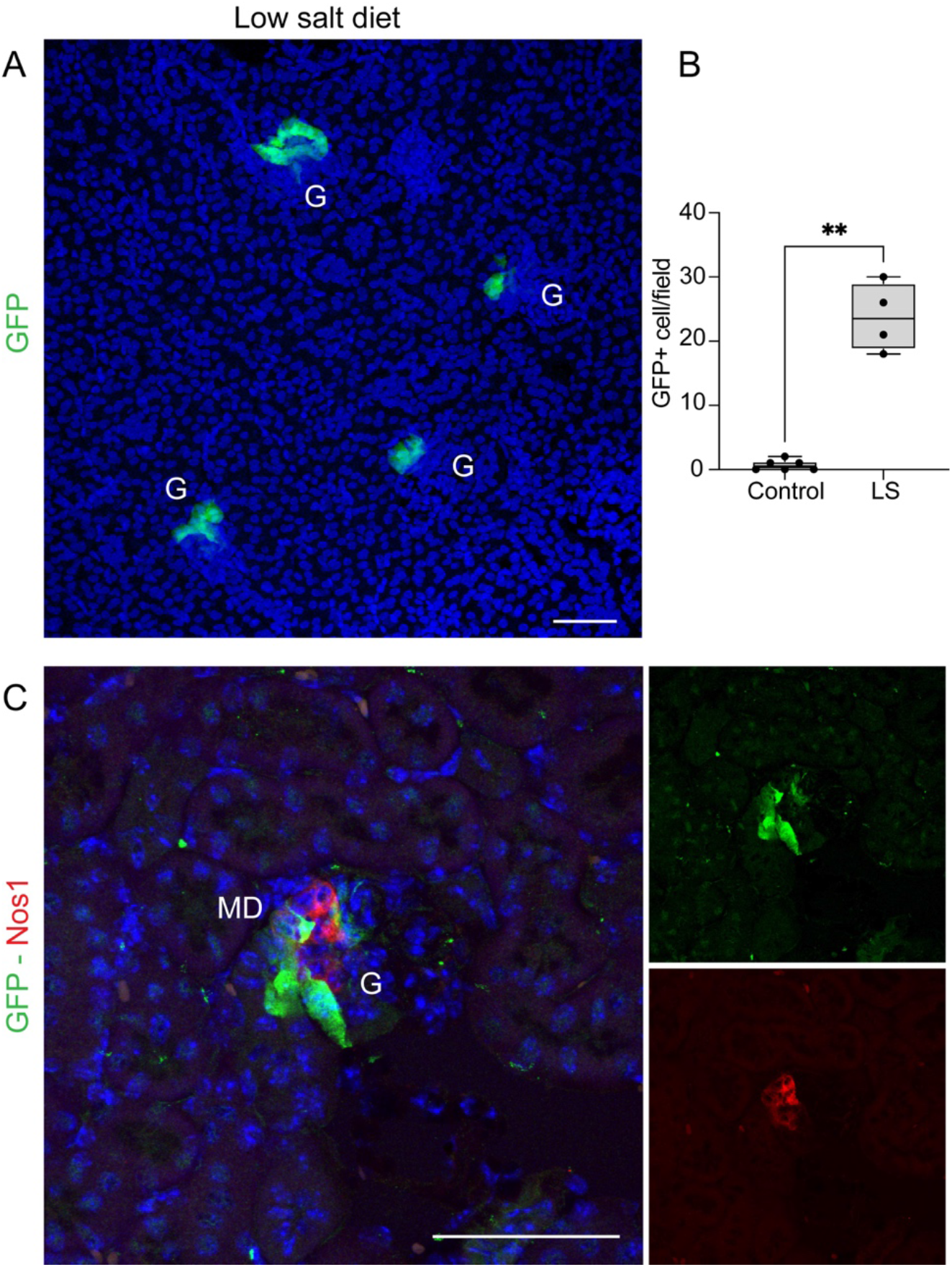
Immunofluorescence labeling of GFP in kidney tissues of AAV9-GFP-injected adult mice on a low-salt (LS) diet. A: Overview GFP immunofluorescence (green) image of a large renal cortical area with multiple glomeruli (G). Cell nuclei were labeled blue with DAPI. Note the presence of several GFP+ cells at the vascular pole area of each glomerulus. **B:** Statistical summary of the number of GFP+ cells per field. Data are presented in box plots as a center line (mean), box (25th–75th percentiles), and whiskers (minimum to maximum values). **: p<0.01 with t-test, n=4-6 mice each; data points represent the average of 5 different tissue areas in each. **C:** Immunofluorescence double labeling of the AAV9-driven genetic expression of GFP (green, top right) and the macula densa (MD) cell marker Nos1 (red, bottom right). The overlay image (left) shows partial co-localization (yellow), confirming GFP expression in MD cells and in a few perimacular cells in the thick ascending limb. Scale bars are 50 µm.

### AAV9-mediated MD cell *Pappa2* and *Ccn1* genetic modifications in vivo

Based on the highly MD-specific tropism of wild-type AAV9 in the neonatal kidney and its ability to deliver the GFP reporter RNA (Figure 1), the next series of experiments aimed to determine whether AAV9 can be used in postnatal mice to perform biologically relevant genetic modifications specifically in MD cells in vivo. Because *Pappa2* (Pregnancy-Associated Plasma Protein A2) is by far the most MD-specific differentially expressed gene (DEG) and *Ccn1* (Cell Communication Network 1) is one of the top secreted factors by MD cells (16), we used *Pappa2* knockdown (loss-of-function, MD-Pappa2^Lof^) and *Ccn1* overexpression (gain-of-function, MD-Ccn1^Gof^) as the first applications, delivering *Pappa2* short hairpin RNA (shRNA) or *Ccn1* mRNA, respectively, as the AAV9 cargo. Single ip. injection of low-dose AAV9-control scrambled RNA or AAV9-Pappa2 shRNA (10^11^ GC in 25 μL saline) was given to neonatal mice on postnatal days P2-3. Compared to control, the treatment with AAV9-Pappa2 shRNA resulted in >90% silencing (knockdown) of MD-specific Pappa2 expression (MD-Pappa2^Lof^) based on RNAscope performed on kidney tissue sections harvested one month after the AAV9 injections (Fig. 3, A and B). On the other hand, the injection of AAV9-Ccn1 mRNA (10^11^ GC in 25 μL saline) in a different set of healthy C57BL6 mice at P2-3 resulted in >5-fold MD-specific Ccn1 overexpression (MD-Ccn1^Gof^) compared to control, as confirmed by RNAscope on histological sections of kidney tissues harvested one month later (Fig. 3, C and D).

**Figure 3.**
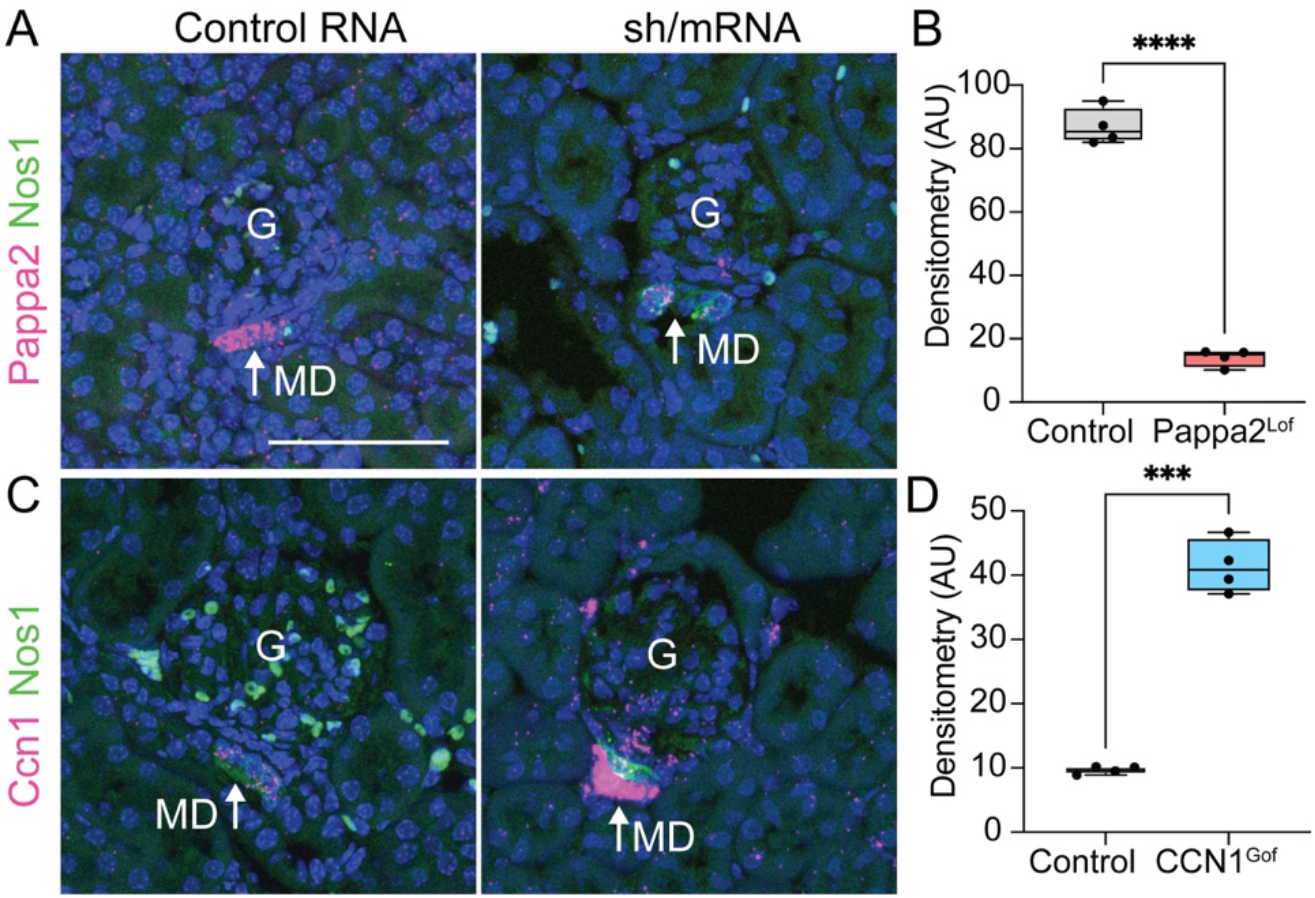
Validation of MD-specific Pappa2 knockdown and Ccn1 overexpression in vivo by AAV9-mediated RNA delivery. A: Representative fluorescence images of double-labeling with the macula densa (MD, arrows) cell markers Nos1 (green, immunofluorescence) and Pappa2 (magenta, RNAscope) in fixed kidney tissue sections from healthy C57BL6 mice treated at postnatal day P2 with a single ip. injection of AAV9-scrambled RNA (control, left) or AAV9-Pappa2 shRNA (right). Nuclei were labeled blue with DAPI. Scale bar is 50 µm for all panels. **B:** Quantification of Pappa2 expression in whole MD plaques based on RNAscope densitometry. Data are presented in box plots as a center line (mean), box (25th–75th percentiles), and whiskers (minimum to maximum values). ****P<0.0001 with t-test, n=4 each. **C:** Representative fluorescence images of double-labeling with the macula densa (MD, arrows) cell markers Nos1 (green, immunofluorescence) and Ccn1 (magenta, RNAscope) in fixed kidney tissue sections from healthy C57BL6 mice treated at postnatal day P2 with a single ip. injection of AAV9-scrambled RNA (control, left) or AAV9-Ccn1 RNA (right). Nuclei were labeled blue with DAPI. **D:** Quantification of Ccn1 expression in whole MD plaques based on RNAscope densitometry. Data are presented in box plots as a center line (mean), box (25th–75th percentiles), and whiskers (minimum to maximum values). ***P<0.001 with t-test, n=4 each.

Functional phenotyping of AAV9-mediated MD cell genetic modifications was performed at the single-cell and single-nephron level using intravital multiphoton microscopy (MPM) imaging of the intact living kidney in wild-type control and MD-Pappa2^lof^ mice one month after the postnatal ip. injection of a single low-dose AAV9-control RNA or AAV9-Pappa2 shRNA (10^11^ GC in 25 μL saline). The MD specificity of AAV9-mediated gene delivery was confirmed by the entirely MD-specific expression of GFP (Fig. 4A). Compared to wild-type controls, MD-Pappa2^lof^ mice featured smaller glomeruli with significantly reduced afferent (Fig. 4B) and efferent arteriole (Fig. 4C) and glomerular diameter (Fig. 4D), glomerular filtration rate (Fig. 4E), diminished magnitude and frequency of tubuloglomerular feedback (TGF) oscillations (Fig. 4, F and G) and reduced single-nephron blood flow (Fig. 4H).

**Figure 4.**
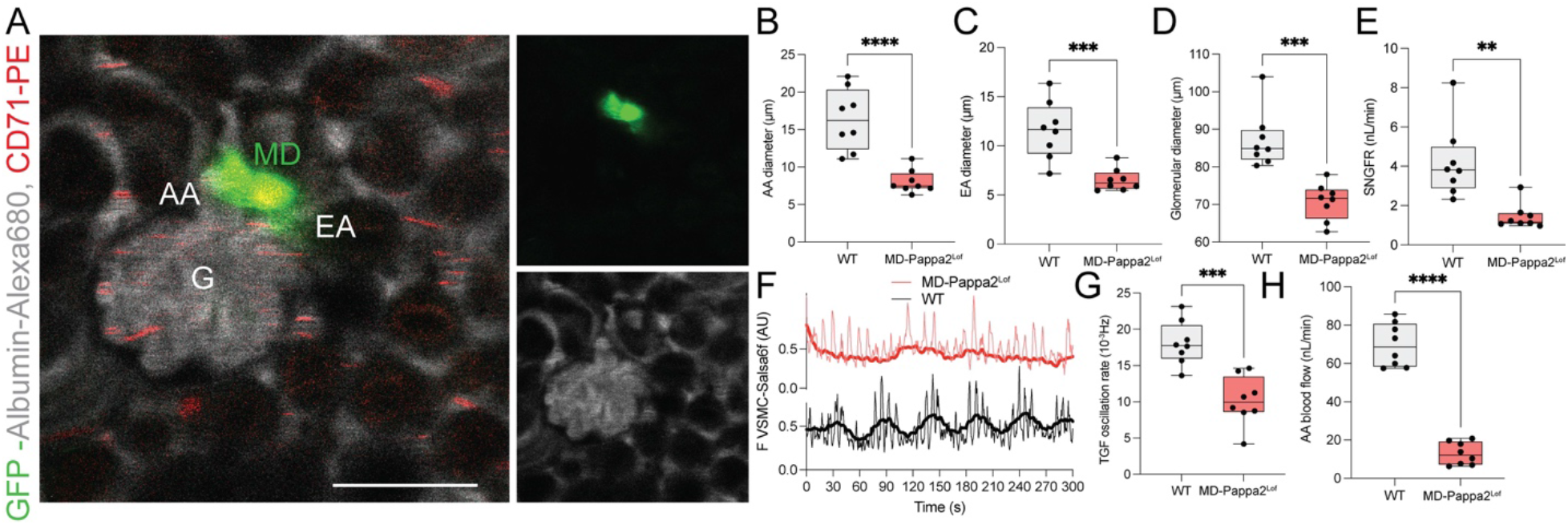
Functional phenotyping of the MD-Pappa2^Lof^ mouse kidneys using intravital MPM imaging. A: Representative in vivo MPM image of a single glomerulus (G) with afferent (AA) and efferent arterioles (EA) and surrounding peritubular capillaries and tubule segments, including the GFP+ macula densa (MD) in a MD-Pappa2^Lof^ mouse kidney. Multi-color labeling of the AAV9-driven genetic expression of GFP (green, top right) and the plasma marker albumin-Alexa Fluor 680 injected iv (gray, bottom right). The overlay image (left) demonstrates MD-specific GFP expression and circulating CD71+ reticulocytes (red) in the glomerular and peritubular capillaries. The scale bar is 50 µm. **B-E:** Statistical summary of the afferent arteriole (AA)(B), efferent arteriole (EA)(C), glomerular diameter (D), and single nephron glomerular filtration rate (SNGFR)(E) in wild-type (WT) control and MD-Pappa2^Lof^ mice. Data are presented in box plots as a center line (mean), box (25th–75th percentiles), and whiskers (minimum to maximum values). **,***,****P<0.01, 0.001, 0.0001 with t-test, n=4 mice each, n=2 nephrons analyzed per mouse. **F:** Time-lapse recordings of the endogenous TGF-mediated oscillations in AA vascular smooth muscle cell (VSMC) calcium (based on Salsa6f fluorescence (F) intensity) in wild-type (WT, black line) control and MD-Pappa2^Lof^ Ren1d-GCaMP6f/tdTomato (Salsa6f) mice (red line). **G-H:** Statistical summary of the rate of TGF oscillations (G) and single nephron (AA) blood flow (H) in wild-type (WT) control and MD-Pappa2^Lof^ mice. Data are presented in box plots as a center line (mean), box (25th–75th percentiles), and whiskers (minimum to maximum values). ***,****P<0.001, 0.0001 with t-test, n=4 mice each, n=2 nephrons analyzed per mouse.

Histological analysis of MD-Pappa2^Lof^ mouse kidneys found smaller glomeruli (based on reduced glomerular tuft area, Fig. 5A, B), reduced number of JG renin cells (based on renin immunofluorescence, Fig. 5C, D) and podocytes (based on WT1 immunofluorescence, Fig. 5E, F) and reduced glomerular capillary density (based on densitometry of CD31 immunofluorescence, Fig. 5G, H) compared to control. In contrast, MD-Ccn1^Gof^ mouse kidneys had larger glomeruli (based on increased glomerular tuft area, Fig. 5A, B) and increased glomerular capillary density (based on CD31 immunofluorescence densitometry, Fig. 5G, H) compared with control.

**Figure 5.**
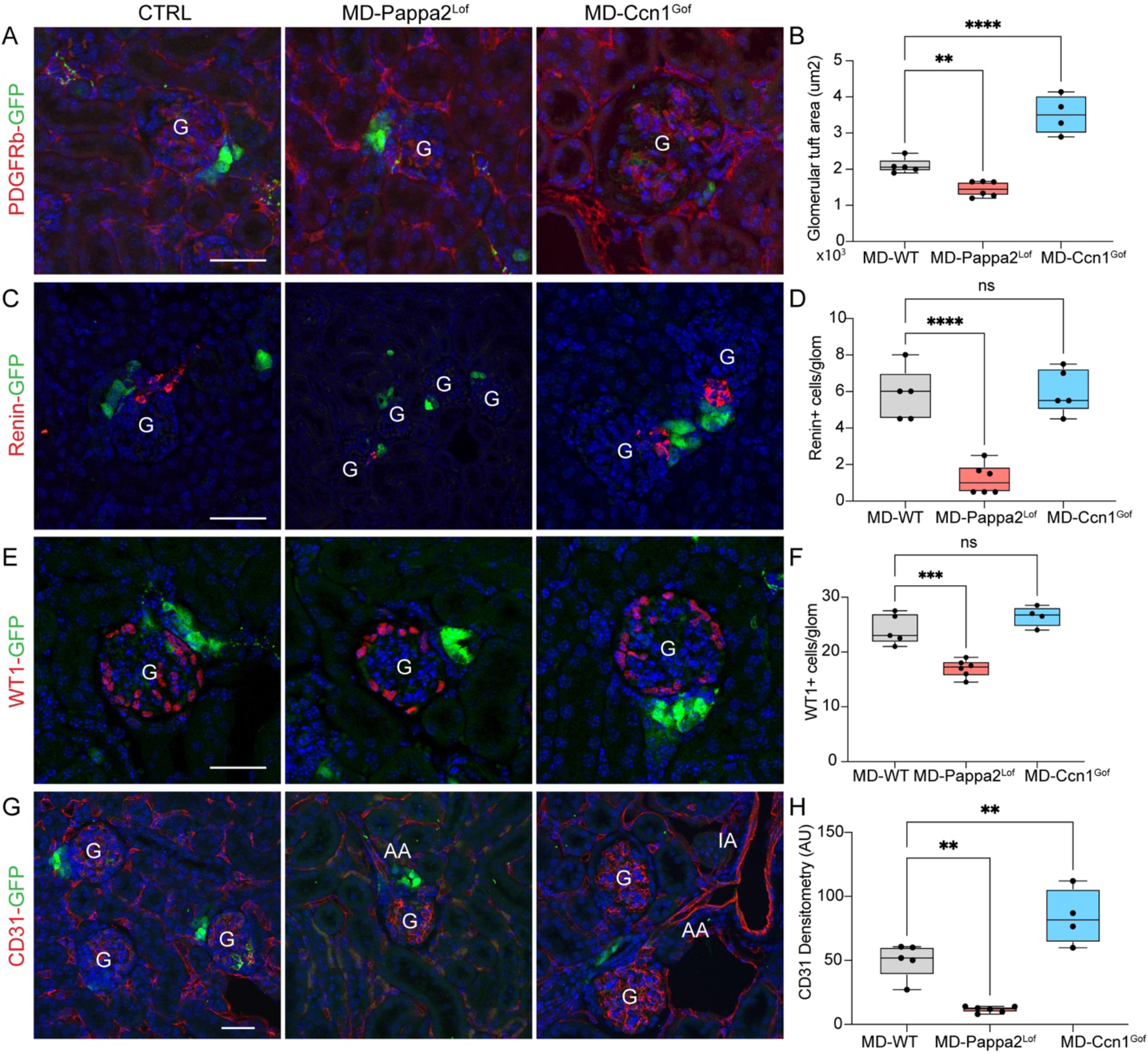
Histological analysis of MD-Pappa2^Lof^ and MD-Ccn1^Gof^ mouse kidneys. A-H: Representative immunofluorescence images with double labeling of the macula densa (MD)-specific, AAV9-driven genetic expression of GFP (green) and either PDGFRb (A), renin (C), WT1 (D), and CD31 (G)(red) in wild-type (WT) control (AAV9-scrambled RNA-treated, left column), MD-Pappa2^Lof^ (AAV9-mPappa2 shRNA-treated, center column), and MD-Ccn1^Gof^ (AAV9-mCcn1 RNA-treated, right column) mouse kidney section. Nuclei were labeled blue with DAPI. G: glomerulus, AA: afferent arteriole, IA: interlobular arteriole. **B, D, F, H:** Statistical summary of the glomerular tuft area (B), the number of renin+ (D) and WT1+ cells (F), and CD31 densitometry (H) per glomerulus. Data are presented in box plots as a center line (mean), box (25th–75th percentiles), and whiskers (minimum to maximum values). Ns: not significant, **,***,****P<0.01, 0.001, 0.0001 with t-test, n=4-6 mice each, n=5 glomeruli analyzed per mouse. Scale bar is 50 μm for all panels.

## Discussion

This study demonstrated the high affinity of wild-type AAV9 for the minority renal cell type of the macula densa in the neonatal stage and after a temporary physiological challenge (low-salt diet), as well as the feasibility of performing MD cell-specific genetic manipulations in the kidney in vivo using this AAV9 approach. After confirming high MD cell specificity and efficiency using AAV9-mediated GFP reporter expression, proof-of-concept studies used AAV9 to overexpress or knock down biologically relevant genes in MD cells and establish MD gain- and loss-of-function mouse models for initial functional and structural phenotyping. RNAscope confirmed the MD-specific knockdown of Pappa2 expression by AAV9-Pappa2 shRNA treatment in MD-Pappa2^Lof^ mice, and the MD-specific overexpression of Ccn1 by AAV9-Ccn1 mRNA treatment in MD-Ccn1^Gof^ mice. The mouse phenotypes revealed alterations in classic MD and JGA functions (renin control and TGF-mediated glomerular hemodynamics) as well as in newly emerging MD cell functions (kidney tissue remodeling and regeneration).

The MD-targeting gene delivery approach described here adds an exciting new component to the many uses of AAVs, including AAV9 for kidney cell targeting genetic manipulations in vivo, as established in prior work (3–6, 29). While these earlier studies demonstrated the important role of AAV serotype and route of administration in renal cell type-specific tropism, the present study highlighted the additional importance of age dependence (neonatal vs adult) and physiological stimulation conditions (control vs low-salt diet) (Figs. 1-2). In addition to proximal tubule cells, mesangial cells, and podocytes, the selective targeting of “perimacular cells”, the few tubular epithelial cells in the thick ascending limb directly adjacent to Nos1+ MD cells has been described earlier (5, 30). However, MD cells themselves were not transduced by these previously used AAV serotypes and approaches (5). Therefore, the present study and approach using wild-type AAV9 in neonatal mice or after a temporary low-salt diet in adults is the first to achieve a high degree of MD cell specificity (Figs. 1-2).

Given the robust MD specificity of AAV9-mediated gene delivery in neonatal animals, this approach allows for rapid, relatively inexpensive generation of chimeric animals for MD targeting gene overexpression (MD gain-of-function models), or gene knock-down (MD loss-of-function models) in vivo, as demonstrated by the first proof-of-concept experiments in this study (Figs. 3-5). The initial results using MD-Pappa2^Lof^ mice confirmed that MD Pappa2 is important for normal glomerular and vascular development and renal tissue maintenance in the adult kidney. The smaller glomeruli with reduced numbers of renin, endothelial cells, and podocytes (Figs. 4-5), and reduced renal and glomerular hemodynamics (Fig. 4) observed in MD-Pappa2^Lof^ mice at the single-nephron level are consistent with the absence of MD cell Pappa2, a locally produced and secreted tissue hypertrophy factor (16, 17, 31). Since the control of renin release and TGF are classic MD cell and JGA functions (11–14), the alterations in renin and TGF oscillations observed in MD-Pappa2^Lof^ mice (Figs. 4-5) further substantiate MD cell specificity of the applied AAV9 gene delivery approach. Considering that Pappa2 is the highest ranked differentially expressed gene in MD cells, these results further confirm the recently reported tissue remodeling and regenerative function of MD cells (15, 16). Future studies are needed to determine the global renal and systemic functions of MD-Pappa2 and the effects of the hypodynamic hemodynamic state (hypoperfusion, hypofiltration) observed in MD-Pappa2^Lof^ mice on physiological functions and in disease states. As another proof-of-concept experiment, MD-Ccn1^Gof^ mice in this study demonstrated the feasibility of generating and studying AAV9-mediated MD-specific gene overexpression (Fig. 3). The larger glomeruli and increased glomerular capillary density observed in MD-Ccn1^Gof^ mice (Fig. 5) confirm the recently reported pro-angiogenic and tissue remodeling functions of MD-derived Ccn1 (15, 16).

Besides the high-degree MD cell specificity and targeting of wild-type AAV9 in the neonatal kidney, the present study found a considerable amount of GFP+ cells in the brain, heart, and liver (Fig. 1E). These results are consistent with earlier work that described AAV9 tropism for the neonatal brain (9) and heart (8), and the generally high affinity of most AAVs for the liver (2). Although the amount of off-target (non-MD) cells in these organs was moderate, likely due to the applied low-dose AAV9 (Fig. 1E), extra-renal targeting by AAV9 may cause complications when interpreting MD-focused functional data at the systemic level. In future work, the combination of this AAV9-mediated gene delivery approach with using MD cell-specific gene promoters in the design of the RNA construct can further increase MD cell-specificity of the genetic manipulations.

Similarly to the findings in brain tissue (9), AAV9-mediated gene delivery to the kidney was inefficient after day P10 in control mice (Fig. 1A). The total lack of AAV9-mediated transduction seen in the adult kidney suggests there is a short developmental period in which JGA access by circulating virus to MD cells becomes restricted. Rapid JGA tissue maturation during the early neonatal period, especially the reduction in JG renin cell number and the shortening of the renin-positive segment of the afferent arteriole towards the JGA (32), which is covered by fenestrated endothelium (33), may determine the time dependence of successful AAV9 transduction of MD cells. Prior work has demonstrated the presence of a fenestrated endothelium, preglomerular plasma ultrafiltration, and bulk fluid flow through the renin-positive segment of the afferent arteriole into the JGA interstitium (33, 34). Reactivation of this afferent arteriole-JGA developmental pattern (increased renin+ segment and JGA access) by low-salt diet may explain the re-established MD cell targeting of AAV9 by this condition in the adult (Fig. 2). In addition to these microanatomical conduits that grant JGA access for circulating AAV9, the high degree of endocytosis and vesicle trafficking (17) and the high expression of cell surface glycans (known AAV9 attachment receptors) in MD cells (16) may further explain the high affinity of AAV9 for MD cells. Given that prior studies established the anterograde transsynaptic spread of AAV9 across neuronal synapses (35), it is important to note that neuronally differentiated MD cells are innervated by both sympathetic efferent and sensory afferent nerves (19), which were also targeted by AAV9 in the present study (Fig. 1G). However, because GFP+ renal nerves were present, but MD cells were not GFP-labeled in the adult (Fig. 1G), we can exclude the possibility that MD cell targeting in our work resulted from transsynaptic spread of AAV9 from renal nerves.

In conclusion, this study demonstrated the high affinity of wild-type AAV9 for MD cells in the neonatal kidney and after a temporary low-salt diet in the adult and established an important new element of the MD cell research toolbox. MD-specific genetic manipulations in vivo via AAV9-mediated gene delivery may open the door to the further characterization of MD cell functions, including the MD program for endogenous kidney tissue remodeling and regeneration, as well as for the development of MD cell-targeting gene therapies.

## Grants

This work was supported by US National Institutes of Health grants R01DK064324, R01DK123564, R01DK135290, and S10OD038245 to J.P-P, and R01DK144444 to G.G.

## Author contributions

G.G. and J.P.P. designed the study and wrote the manuscript. A.B.C., A.I., V.B., S.D., and K.A. made substantial contributions to data acquisition. G.G., A.B.C., A.I., and J.P.P. analyzed the imaging data. All authors approved the final version of the manuscript.

## Acknowledgments

Fluorescence imaging was performed at the USC Multi-Photon Microscopy Core using a Leica Stellaris 8 Falcon multimodal multiphoton microscope with a Chameleon Discovery NX laser, purchased with the NIH shared instrument grant 1S10OD038245.

